# rMETL: sensitive mobile element insertion detection with long read realignment

**DOI:** 10.1101/421560

**Authors:** Tao Jiang, Bo Liu, Yadong Wang

## Abstract

**Summary:** Mobile element insertion (MEI) is a major category of structure variations (SVs). The rapid development of long read sequencing provides the opportunity to sensitively discover MEIs. However, the signals of MEIs implied by noisy long reads are highly complex, due to the repetitiveness of mobile elements as well as the serious sequencing errors. Herein, we propose Realignment-based Mobile Element insertion detection Tool for Long read (rMETL). rMETL takes advantage of its novel chimeric read re-alignment approach to well handle complex MEI signals. Benchmarking results on simulated and real datasets demonstrated that rMETL has the ability to more sensitivity discover MEIs as well as prevent false positives. It is suited to produce high quality MEI callsets in many genomics studies.

**Availability and Implementation:** rMETL is available from https://github.com/hitbc/rMETL.

**Supplementary information**: Supplementary data are available at *Bioinformatics* online.

## 1 Introduction

Mobile element insertion (MEI) represents about 25% of structure variations (SVs) in human genome, which are principally contributed by active mobile elements, such as Alu, L1 and SVA families (Stewart *et al*., 2011). Efforts have been made to detect MEIs with short reads (Gardner *et al*., 2017), however, short read-based approaches could have limitations when dealing with highly repetitive mobile elements.

Long reads have demonstrated their better ability to handle repeats and discover SVs (Sedlazeck et al, 2018a). However, with both of the repetitiveness of mobile elements and serious sequencing errors, the signals of MEI implied by long reads are highly complex (a detailed discussion is in Supplementary Notes). State-of-the-art long read-based SV detection approaches (Sedlazeck et al, 2018b) use unified approaches to detect various kinds of SVs. This “one-fits-all” strategy may be not able to fully consider the characteristics of MEI to well handle the complex signals.

Herein, we propose Realignment-based Mobile Element insertion detection Tool for Long read (rMETL). rMETL takes advantage of its specifically designed chimeric read re-alignment approach to well handle the complex MEI signals to sensitively discover MEIs as well as prevent false positives. Benchmarking results demonstrate that rMETL can produce high quality callsets to improve long read-based MEI calling.

## 2 Methods

rMETL supports the reads produced by mainstream platforms such as Pacific Biosciences (PacBio) and Oxford Nanopore Technology platforms. Using sorted BAM files as input, rMETL extracts and re-aligns chimerically aligned long reads to discover MEIs in four steps as following (schematic illustrations are in Supplementary Figs.1 and 2).

1) rMETL extracts chimeric reads (reads having split alignment, large clippings or large indels) from input files (Supplementary Fig. 2);

2) rMETL clusters the chimerically aligned read parts with a set of specifically designed rules, to infer a set of infers putative MEI sites;

3) rMETL realigns the clustered read parts to the consensus sequences of Alu, L1 and SVA families with a well-tuned read alignment approach;

4) rMETL investigates the realignment results to find out the evidences to call MEIs as well as filter false positive candidates.

Refer to Supplementary Notes for more detailed information.

## 3 Results

We implemented rMETL on simulated and real datasets to assess its ability of MEI calling. A state-of-the-art long read-based SV calling approach, Sniffles (Sedlazeck *et al*, 2018b), is employed for comparison. The details of the implementation of the benchmarking is in Supplementary Notes.

Four PacBio-like simulated datasets (mean read length: 8000 bp) in various sequencing depths (5X, 10X, 20X and 50X, respectively) are generated with an *in silico* donor human genome having 20,000 MEIs to make a baseline assessment. Both of rMETL and Sniffles achieved improving sensitivities with the increase of depths (Table 1 and Supplementary Table 1), moreover, the sensitivity of rMETL is higher especially for the lower depth datasets (5X and 10X), where outperforms Sniffles 10-20%. Moreover, both of rMETL and Sniffles have few false positives.

**Table 1.** sensitivities of the callsets of rMETL and Sniffles on four simulated PacBio datasets in various sequencing depths.

|  | 5X | 10X | 20X | 50X |
| --- | --- | --- | --- | --- |
| rMETL | 49.24% | 78.64% | 88.19% | 93.79% |
| Sniffles | 28.06% | 68.93% | 86.19% | 91.55% |

To further access the accuracy of rMETL, we generated another 50X PacBio-like simulated dataset from an *in silico* donor human genome having 20,000 insertions which none of the inserted sequences are mobile elements, but their lengths are similar to that of Alu, SVA or L1. On this dataset, rMETL make 366 calls, i.e., only 1.8% of the 20,000 events are false positively recognized as MEIs, suggesting that rMETL have good ability to prevent false positives.

There are still a few cases that rMETL cannot work well, mainly due to abnormal read coverages and serious sequencing errors. A discussion is in Supplementary Notes.

We further assess rMETL with a 55X PacBio dataset (Zook *et al*., 2014) and a 28X ONT dataset (Jain *et al*., 2018) from the well-studied CEPH sample NA12878. Due to lack of grand truth, four high quality short read-based MEI callsets (Gardner *et al*., 2017; Lee *et al*., 2012; Thung *et al*.,2014 and Wu *et al*., 2014) are used as pseudo grand truth callsets.

rMETL respectively called 6022 and 5439 MEIs on the PacBio and ONT datasets (Supplementary Table 2). It is observed from Venn diagrams (Fig.1 and Supplementary Fig.3) that, the callsets of rMETL can cover most of the MEIs which are supported by at least two short read-based callsets. Considering that the calls supported by two or more independent approaches have relatively high confidence to be true positive MEIs, this result suggest that rMETL may have high sensitivities. Moreover, it is also worthnoting that the callsets of rMETL covered most of MEIs in the 1000 Genomes Project callset (Sudmant *et al*., 2015).

**Fig 1.**
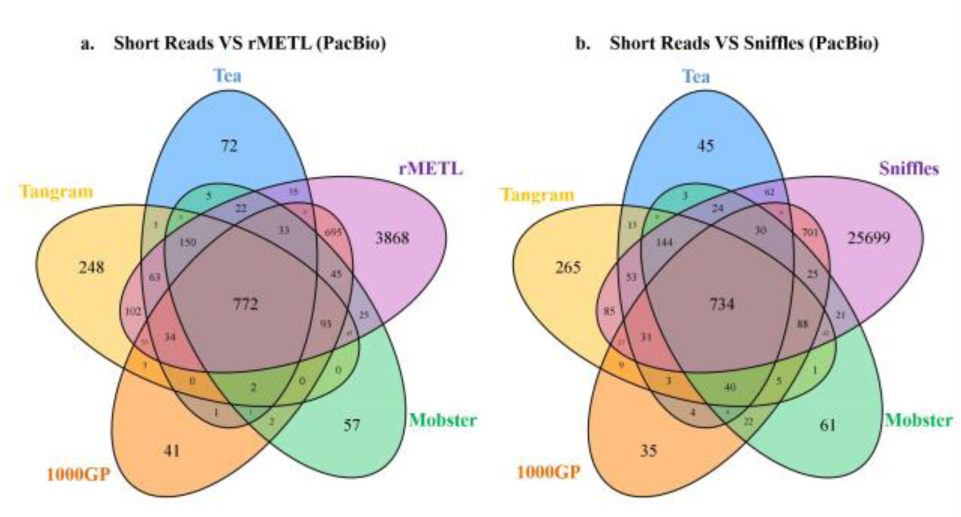
Venn diagrams of the four short read-based MEI callsets of NA12878 and the long read-based callsets respectively made by (a) rMETL and (b) Sniffles.

Sniffles respectively called 27772 and 63925 INS/DELs for the PacBio and ONT datasets. The high numbers of calls are also reasonable since Sniffles calls not only MEIs, but also other kinds of large insertions and deletions. However, for both of the two datasets, the corresponding callsets of Sniffles covered about 5%-6% less MEIs which have at least two short read-based callset supports than that of rMETL (Fig.1, Supplementary Fig.3 and Supplementary Table 2), indicating that the sensitivity of Sniffles could be lower. This is mainly due to that, with the generic design, Sniffles could have relatively poor ability to deal with the complex MEI signals. However, rMETL has its own advantage to use read realignment approach to transform the ambiguous and chimeric read alignments into more homogenous alignments, which produces strong evidences (Supplementary Notes and Supplementary Fig.4).

There are a proportion of MEI calls made by rMETL not supported by any of short read-based callsets (i.e., 3864 and 3287 calls for the PacBio and ONT datasets, respectively, Supplementary Table 2). However, this may not indicate serious false positives. It is observed that, 96% (PacBio) and 87% (ONT) of such calls are also in the callsets of Sniffles (Supplementary Fig.3c-d). This indicates that most of these MEIs could be also plausible. We further investigated these short read-unsupported calls of rMETL, and found that most of them have strong MEI evidences, i.e., there are many chimeric read parts in local regions which can be confidently aligned to the consensus sequences of mobile elements (Supplementary Notes and Supplementary Fig.5).

The runtimes and memory footprints were also assessed (Supplementary Table 3). For the PacBio and the ONT datasets, rMETL can accomplish the task in 2-3 hours with 8 CPU threads and 10-12 GB RAM, which is about 2 and 1.5 times faster than Sniffles, respectively.

Overall, the benchmarking results suggest that rMETL has its own power to more sensitively discover MEIs. Due to the complexity of SVs, it could be non-trivial to well handle all kinds of SVs with a generic approach. We believe that rMETL is suited to be integrated into many pipelines to play important roles in cutting-edge genomics studies. Please also refer to Supplementary Notes for more detailed discussions.

